# Delineation of the pathogenic presynaptic mechanisms of synaptotagmin-1 variants

**DOI:** 10.1101/2023.10.29.564558

**Authors:** Holly Melland, Kyla Venter, Stephanie L. Leech, Amy Ferreira, Elyas H. Arvell, Kate Baker, Sarah L. Gordon

## Abstract

Missense variants in the presynaptic calcium sensor synaptotagmin-1 (SYT1) are associated with an autosomal dominant neurodevelopmental disorder (NDD). These cause dominant-negative impairment of evoked neurotransmitter release in a mutation-specific manner; however, whether NDD-associated variants also perturb the auxiliary functions of SYT1 remains unknown. We investigated whether the expression of SYT1 variants in cultured hippocampal neurons altered either action potential-independent spontaneous synaptic vesicle exocytosis or synaptic vesicle endocytosis. SYT1 variants did not induce dominant-negative impairment of either process, confirming that defective evoked exocytosis is the major pathogenic mechanism of SYT1-associated NDD. To examine the differential impacts of human variants on evoked exocytosis, both NDD-associated and strategic mutations were used to explore the functional importance of two distinct Ca^2+^-binding residues in the C2B domain. We show that both the nature of the amino acid change and the specific residues targeted determine the severity of exocytic disturbance. Together, this work informs understanding of SYT1 function and further clarifies potential mechanistic targets for treating SYT1-associated NDD.

## INTRODUCTION

Synaptotagmin-1 (SYT1) is an essential synaptic vesicle protein that acts as the major Ca^2+^-sensor mediating fast, synchronous neurotransmitter release in the mammalian cerebrum (Bowers and Reist, 2020; Geppert et al., 1994; Marqueze et al., 1995; Maximov and Sudhof, 2005; Rizo, 2022; Ullrich and Sudhof, 1995; Xu et al., 2007). Heterozygous variants in SYT1 give rise to a neurodevelopmental disorder (MIM 618218; Baker-Gordon Syndrome or *SYT1*-associated neurodevelopmental disorder) with 16 published *de novo* variants linked to the disorder to date (Cornelisse et al., 2023; Melland et al., 2022). Affected individuals present with a spectrum of features including motor delay, intellectual disability, behavioural disturbances, and movement disorders, with abnormal electroencephalograms being an additional hallmark (Baker et al., 2018; Melland et al., 2022).

SYT1 is an integral synaptic vesicle protein with two cytoplasmic Ca^2+^-binding C2 domains (C2A and C2B)(Perin et al., 1991). The tertiary structure of each C2 domain includes three loops that together form a pocket where three (C2A) or two (C2B) Ca^2+^ ions are coordinated by five key aspartate residues (termed D1 through D5)(Fernandez et al., 2001; Perin et al., 1991; Sutton et al., 1995). When neurons are at rest, with low presynaptic Ca^2+^ concentration, SYT1 “clamps” the fusion of vesicles with the plasma membrane and thus inhibits spontaneous neurotransmitter release. Following action potential-evoked Ca^2+^ influx, Ca^2+^-binding neutralises the charge of the loops, relieving the clamping action of SYT1 and allowing the loops to penetrate the negatively-charged plasma membrane, led by hydrophobic residues situated at the tips of these loops (Bowers and Reist, 2020; Brunger et al., 2018; Rizo, 2022). SYT1 also binds to the soluble N-ethylmaleimide-sensitive factor attachment protein receptor (SNARE) complex (Zhou et al., 2015; Zhou et al., 2017). In cooperation with the force provided by zippering SNARE complexes, membrane penetration by SYT1 facilitates fusion of synaptic vesicles with the presynaptic plasma membrane and the release of neurotransmitters (Bowers and Reist, 2020; Brunger et al., 2019; Rizo, 2022).

Loss of SYT1 dramatically reduces action potential-evoked exocytosis and desynchronises residual neurotransmitter release from the stimulus (Courtney et al., 2019; Geppert et al., 1994; Maximov and Sudhof, 2005; Nishiki and Augustine, 2004b). Four pathogenic *SYT1* variants have been demonstrated to impair action potential-evoked neurotransmitter release in a dominant-negative manner (Baker et al., 2015; Baker et al., 2018; Bradberry et al., 2020). These variants, equating to D303G, D365E, I367T and N370K in the rat SYT1 sequence (which is used for simplicity throughout this paper), are situated in the Ca^2+^-binding and membrane-penetrating loops of the C2B domain (Baker et al., 2018). D303 and D365 are key Ca^2+^-binding residues (D1 and D4, respectively), I367 is a hydrophobic tip residue important for membrane penetration, and N370 may aid stabilisation of loop 3 (Baker et al., 2015; Fernandez et al., 2001).

SYT1 also regulates other aspects of synaptic physiology, including modulating the endocytic recycling of synaptic vesicles (Poskanzer et al., 2006; Poskanzer et al., 2003; Yao et al., 2012) in addition to suppressing, or “clamping”, spontaneous exocytosis of synaptic vesicles in the absence of activity (Courtney et al., 2019; Vevea and Chapman, 2020; Xu et al., 2009). A key unresolved question is whether, in addition to impairing evoked neurotransmitter release, pathogenic SYT1 variants also impact these auxiliary functions of SYT1 in a dominant-negative manner. Absence, depletion, or cleavage of SYT1 results in an increase in spontaneous neurotransmitter release (Courtney et al., 2019; Kerr et al., 2008; Pang et al., 2006; Vevea and Chapman, 2020; Xu et al., 2009). Additionally, some, but not all, mutations in SYT1 that inhibit evoked release also increase spontaneous exocytic events in mammalian neurons ((Rhee et al., 2005; Xu et al., 2009; Zhou et al., 2015), but see (Brewer et al., 2015; Huson et al., 2019; Li et al., 2006; Wu et al., 2022; Xu et al., 2009; Zhou et al., 2015)). It was shown that one disease-associated SYT1 variant (D303G) failed to fully restore clamping of spontaneous release when expressed in SYT1 knockout (KO) neurons, though two other pathogenic variants (D365E and I367T) were able to at least partially rescue this clamping function (Bradberry et al., 2020). However, the effect of these variants on spontaneous exocytosis has never been examined in a model of the heterozygous disease state.

Furthermore, loss of SYT1 disrupts endocytosis of synaptic vesicles (Li et al., 2017; Nicholson-Tomishima and Ryan, 2004; Poskanzer et al., 2003; Yao et al., 2011; Yao et al., 2012). The specific motifs in SYT1 that mediate this modulation of endocytosis have not been definitively mapped, though the Ca^2+^-binding ability of SYT1 has been implicated (Chen et al., 2022; Poskanzer et al., 2006; Yao et al., 2011). Previous studies have suggested that disorder-associated SYT1 variants may disrupt endocytosis or the endocytic trafficking of SYT1 (Baker et al., 2015; Baker et al., 2018), but effects on the kinetics of endocytosis have not been assessed for all variants.

Additionally, the factors contributing to the variation in neurodevelopmental phenotypes observed in *SYT1*-associated neurodevelopmental disorder remain unclear. The recurrent D365E variant is associated with a milder clinical phenotype (e.g. less severe intellectual disability, no movement disorder, less delayed motor milestones) and milder impairment of exocytosis than the other variants, suggesting a genotype-phenotype relationship in this disorder (Baker et al., 2018; Bradberry et al., 2020; Melland et al., 2022). However, it is unknown whether the milder functional impact and associated phenotype is driven exclusively by a more conservative substitution, or whether the D365 (D4) locus is less important than D303 (D1) for triggering of exocytosis.

This work therefore extends the functional assessment of disease-associated SYT1 C2B-domain variants by determining whether perturbation of other aspects of synaptic vesicle dynamics, beyond evoked exocytosis, contributes to pathogenesis. Moreover, we explore the molecular mechanisms defining the variable impact of SYT1 variants on the severity of exocytic impairment, which may contribute to divergent clinical phenotypes.

## RESULTS

### SYT1 variants are correctly localised to synaptic vesicles

Although pathogenic SYT1 variants are enriched at presynaptic terminals at a similar level to the WT protein (Baker et al., 2015; Baker et al., 2018; Bradberry et al., 2020) it is imperative to also establish whether these variants might impact the targeting of SYT1 to synaptic vesicles before functional assays can be accurately interpreted. Quantification of vesicular SYT1 and measurement of synaptic vesicle dynamics were performed using a pH-sensitive fluorescent reporter, pHluorin, conjugated to the lumenal domain of SYT1 (SYT1-pHluorin). The pHluorin moiety fluoresces upon exposure to the neutral extracellular environment during exocytosis and is then quenched upon the reacidification of synaptic vesicles following endocytosis (Miesenbock et al., 1998; Sankaranarayanan et al., 2000). SYT1- pHluorin, either harbouring pathogenic SYT1 variants or the WT SYT1 protein, was transfected into hippocampal neuronal cultures. SYT1-pHluorin situated on the cell surface is fluorescent at rest, allowing plasma membrane-localised SYT1-pHluorin to be quantified as a percentage of total SYT1- pHluorin in the presynaptic terminal (Fig 1A). The difference between total SYT1-pHluorin and cell surface SYT1-pHluorin yields the percentage of internalised SYT1-pHluorin resident on synaptic vesicles within the nerve terminal. Partitioning of SYT1-pHluorin between synaptic vesicle and plasma membranes was similar to that of WT SYT1 for all SYT1 variants (Figure 1B, Table S1), confirming that these mutations do not disturb protein trafficking. The pathogenic exocytic defects caused by SYT1 variants are therefore not mediated by mislocalisation of SYT1 to the plasma membrane (and concomitant reduction in vesicular SYT1).

**Figure 1.**
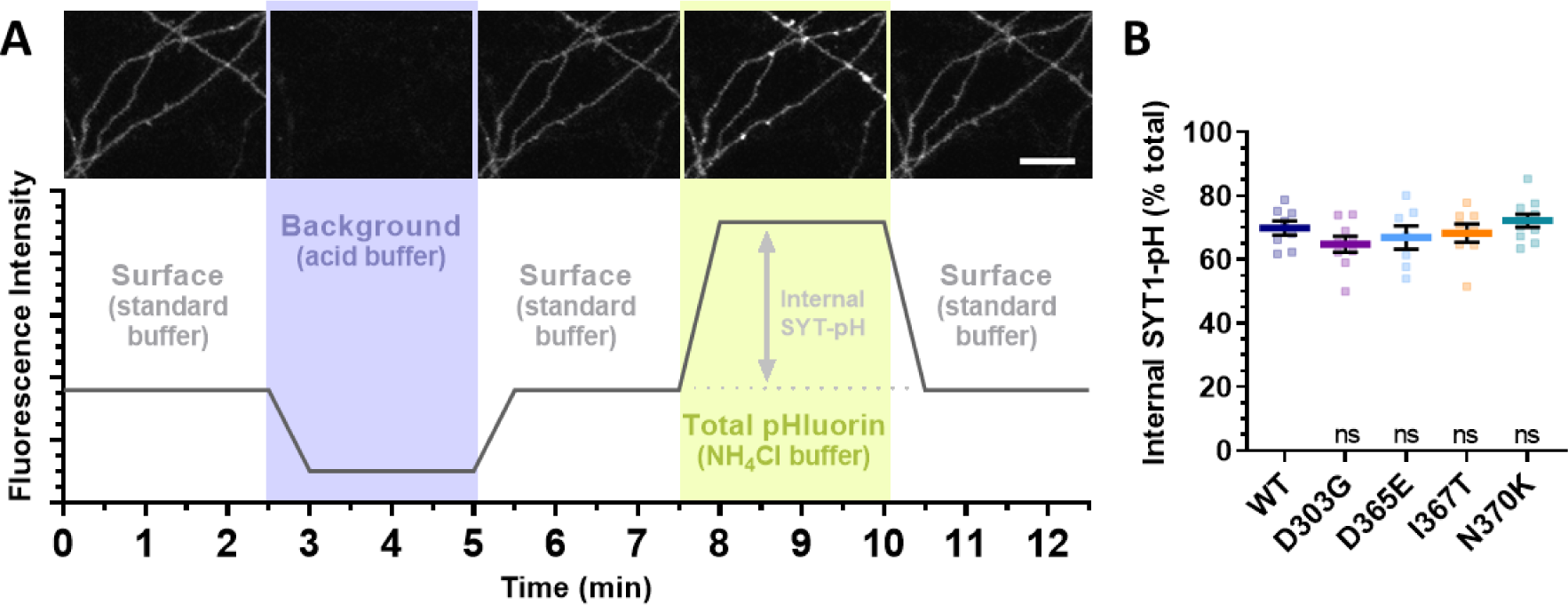
SYT1 variants are correctly localised to synaptic vesicles. Neurons were transfected with SYT1-pHluorin variants (WT, D303G, D365E, I367T or N370K), and cultures were imaged at 37°C while successively superfused with standard, acidic, and NH_4_Cl buffers. A) Top: Representative images (cropped field of view) of SYT1-pHluorin fluorescence corresponding to the experimental paradigm below. Scale bar represents 50 μm. Bottom: Experimental paradigm for assessing the internal fraction of SYT1-pH variants. B) To quantify the localisation of SYT1-pHluorin variants to synaptic vesicles, SYT1- pHluorin fluorescence was normalised to acid buffer (minimum) and NH_4_Cl buffer (maximum) responses and the internal portion (i.e. not on the cell surface) of SYT1-pH was calculated as a percentage of total SYT1-pHluorin in the presynaptic bouton. Values are mean ± SEM, n = 7-10. One-way ANOVA with Dunnett’s multiple comparison’s test was not significant, p>0.05 for all variants compared to WT (see Table S1 for exact P-values). SYT-pH: SYT-pHluorin.

### Pathogenic SYT1 variants do not exert dominant-negative effects on spontaneous synaptic vesicle exocytosis

The impact of pathogenic Ca^2+^-binding loop SYT1 variants on spontaneous release has not been assessed in a heterozygous disease-relevant model system. We probed whether aberrant spontaneous release was a contributing factor to the pathogenesis of SYT1-associated neurodevelopmental disorder by assaying neurons expressing both endogenous WT SYT1 and exogenous mutant SYT1 (in an approximately 1:1 ratio (Baker et al., 2018)). Cultured WT neurons transfected with SYT1-pHluorin variants were assayed in the presence of vATPase-inhibitor bafilomycin, which prevents vesicle re-acidification, so that pHluorin fluorescence reports only exocytic events in a cumulative manner. Cultures were imaged at rest for 10 minutes (with AP5 and CNQX to inhibit network activity) to monitor spontaneous exocytosis over this period (Ramirez et al., 2012; Smillie et al., 2013; Subrahmanyam et al., 2021). Neurons were subsequently stimulated at 10Hz for 30s to elicit action potential-evoked exocytosis, and only “active” nerve terminals capable of evoked exocytosis were included in analysis of spontaneous activity. Finally, neurons were superfused with NH_4_Cl buffer to reveal total SYT1-pHluorin fluorescence, to which data were normalised (Figure 2A). SYT1-pHluorin fluorescence time traces of spontaneous exocytosis were no different between mutant and WT SYT1-transfected synapses (Figure 2A, Table S1). Importantly, the cumulative amount of spontaneous fusion after 10 minutes was similar between all variants and WT SYT1-pHluorin (Figure 2B, Table S1). Moreover, the initial linear rate of spontaneous synaptic vesicle fusion did not differ between neurons expressing WT or any SYT1 variant (Figure 2C, Table S1).

**Figure 2.**
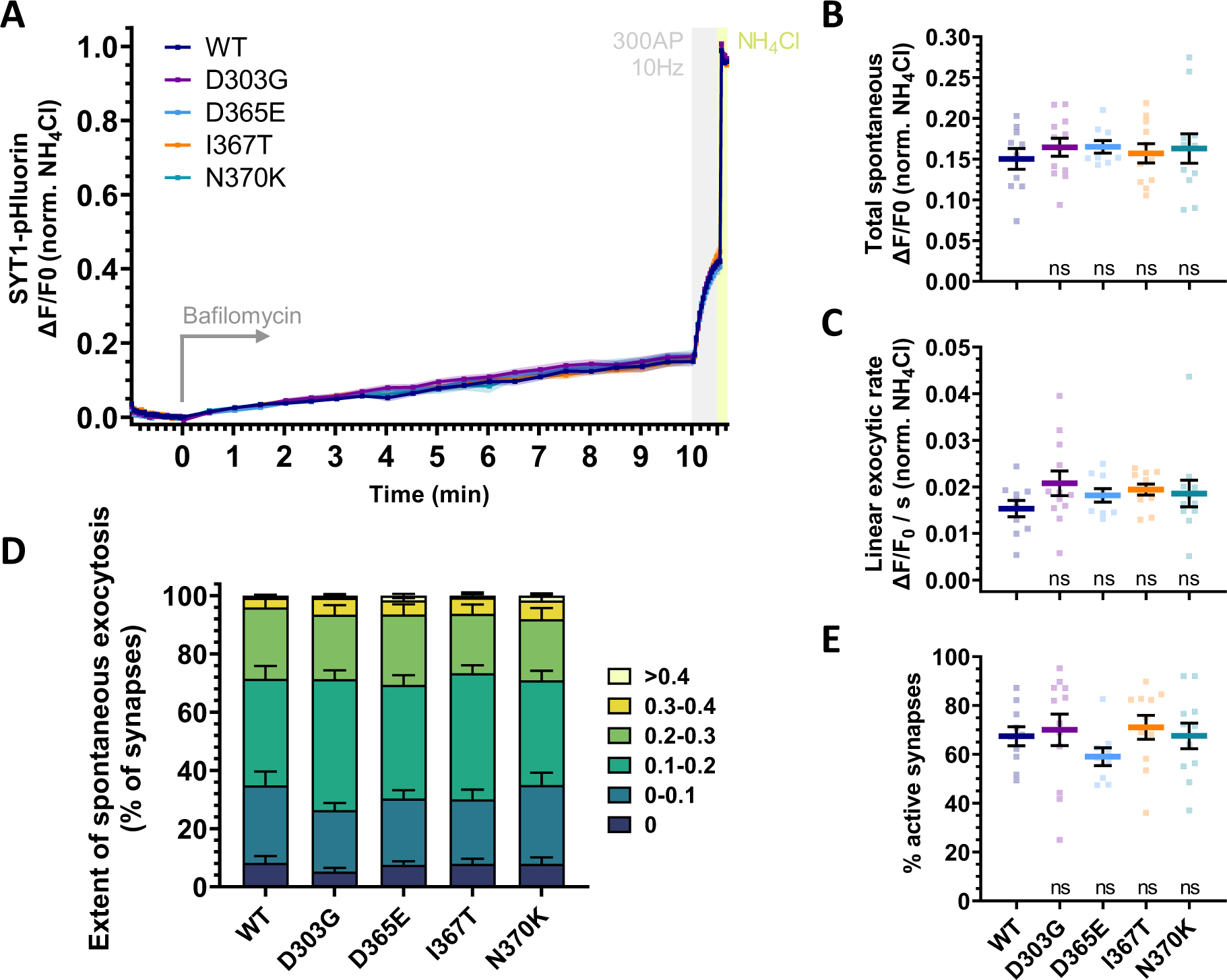
Pathogenic SYT1 variants do not exert dominant-negative effects on spontaneous synaptic vesicle exocytosis. WT neurons were transfected with SYT1-pHluorin variants (WT, D303G, D365E, I367T or N370K) and imaged at 37°C in the presence of 10 μM CNQX and 50 μM AP5. Baseline images were taken for one minute, before 1μM bafilomycin A1 was added and neurons were imaged for 10 minutes at rest to cumulatively measure spontaneous exocytosis. Neurons were then stimulated at 10 Hz for 30 seconds (300 AP) to elicit evoked exocytosis. Neurons were then superfused with NH4Cl buffer to reveal total SYT1-pHluorin fluorescence. Only synaptic puncta responding to stimulation were included in analysis. Fluorescence change from baseline (ΔF/F0) was normalised to the peak of NH4Cl fluorescence. Values are mean ± SEM, n = 9-12. A) Time trace of SYT1-pHluorin fluorescence over the course of the experiment. Two-way repeated measures ANOVA with Dunnett’s multiple comparison’s test was not significant, p>0.05 for all variants compared to WT at all time points. B) SYT1-pHluorin fluorescence (normalised to total NH_4_Cl fluorescence) after 10 minutes of superfusion with bafilomycin, reporting total cumulative spontaneous exocytosis. One-way ANOVA with Dunnett’s multiple comparison’s test was not significant, p>0.05 for all variants compared to WT. C) Linear rate of SYT1-pHluorin fluorescence increase per minute over the first 3 minutes of spontaneous exocytosis. One-way ANOVA with Dunnett’s multiple comparison’s test was not significant, p>0.05 for all variants compared to WT. D) Total amount of spontaneous exocytosis at individual presynaptic boutons (ΔF/F0) normalised to total NH_4_Cl fluorescence, represented as the percent of active synapses in each field of view falling within each bin of total spontaneous exocytosis. Two-way mixed model ANOVA with Dunnett’s multiple comparison’s test was not significant, p>0.05 for all variants compared to WT for each bin. E) Percent of NH_4_Cl-responsive presynaptic boutons in each field of view that showed an increase in SYT1-pHluorin fluorescence upon electrical stimulation, indicating an active synapse. One-way ANOVA with Dunnett’s multiple comparison’s test was not significant, p>0.05 for all variants compared to WT. Exact P-values can be found in Table S1.

There is also the potential for SYT1 variants to alter the number of synapses engaging in substantial spontaneous release. Our pHluorin imaging approach allowed the effect of SYT1 variants to be monitored at the single synapse level. When binning the degree of spontaneous exocytosis at each individual synapse in a field of view, the percentage of synapses falling into each bin was similar for WT SYT1 and all variants (Figure 2D, Table S1). SYT1 variants therefore do not exert notable dominant-negative effects on spontaneous exocytosis, and impaired evoked exocytosis is not accompanied by unclamping of spontaneous exocytosis in SYT1-associated neurodevelopmental disorder. Furthermore, from this assay we could extract the percentage of all presynaptic boutons in each field of view that were active, indicated by exocytosis-induced fluorescence increase in response to electrical stimulation. We observed no difference in the number of active synapses between neuronal cultures transfected with pathogenic variants and WT SYT1-pHluorin (Figure 2E, Table S1).

### Synaptic vesicle endocytosis is not altered by pathogenic SYT1 variants

There is a growing understanding that endocytosis is temperature sensitive and that different stimuli couple to different modes of endocytosis (Chanaday et al., 2019; Chanaday and Kavalali, 2018; Gan and Watanabe, 2018; Lou, 2018; Watanabe and Boucrot, 2017). As such, we sought to examine the impact of pathogenic variants on the kinetics of endocytosis at physiological temperature and with stimuli intended to evoke multiple modes of compensatory endocytosis. Neurons transfected with SYT-pHluorin variants were imaged at 37°C and subjected to one of two different stimulation paradigms: 400AP at 40Hz for 10s (Figure 3A) and 300AP at 10Hz for 30s (Figure 3D). The decay of pHluorin fluorescence following stimulation measures endocytosis of synaptic vesicles. Normalised to the peak fluorescence during stimulation, the time traces of fluorescence decay were no different between WT and mutant SYT1-transfected cultures for either stimulation paradigm (Figures 3A and 3D, Table S1), except for a single timepoint at which D303G was significantly different from WT (P=0.048) following 10Hz stimulation for 30s (Figure 3D, Table S1). This lack of impact of SYT1 variants on endocytosis was corroborated by the endocytic time constants (tau) of post-stimulation fluorescence decay, which were similar to that of WT SYT1-pHluorin for all variants in response to both stimulation paradigms (Figures 3B and 3E, Table S1). Time traces of SYT1-pHluorin fluorescence also reflected the established impact of these variants on evoked exocytosis by exhibiting significant differences from WT during the stimulation phase for I367T and N370K under 40Hz stimulation (Figure 3A, Table S1) and for D303G, D365E, and I367T under 10Hz stimulation (Figure 3D, Table S1). However, the exocytic defects induced by these variants did not significantly alter the fluorescence peak amplitude of response to either stimulation paradigm (Figures 3C and 3F, Table S1). Together, these data from multiple stimulation conditions demonstrate that SYT1 variants do not induce dominant-negative alterations to endocytic rate at physiological temperature.

**Figure 3.**
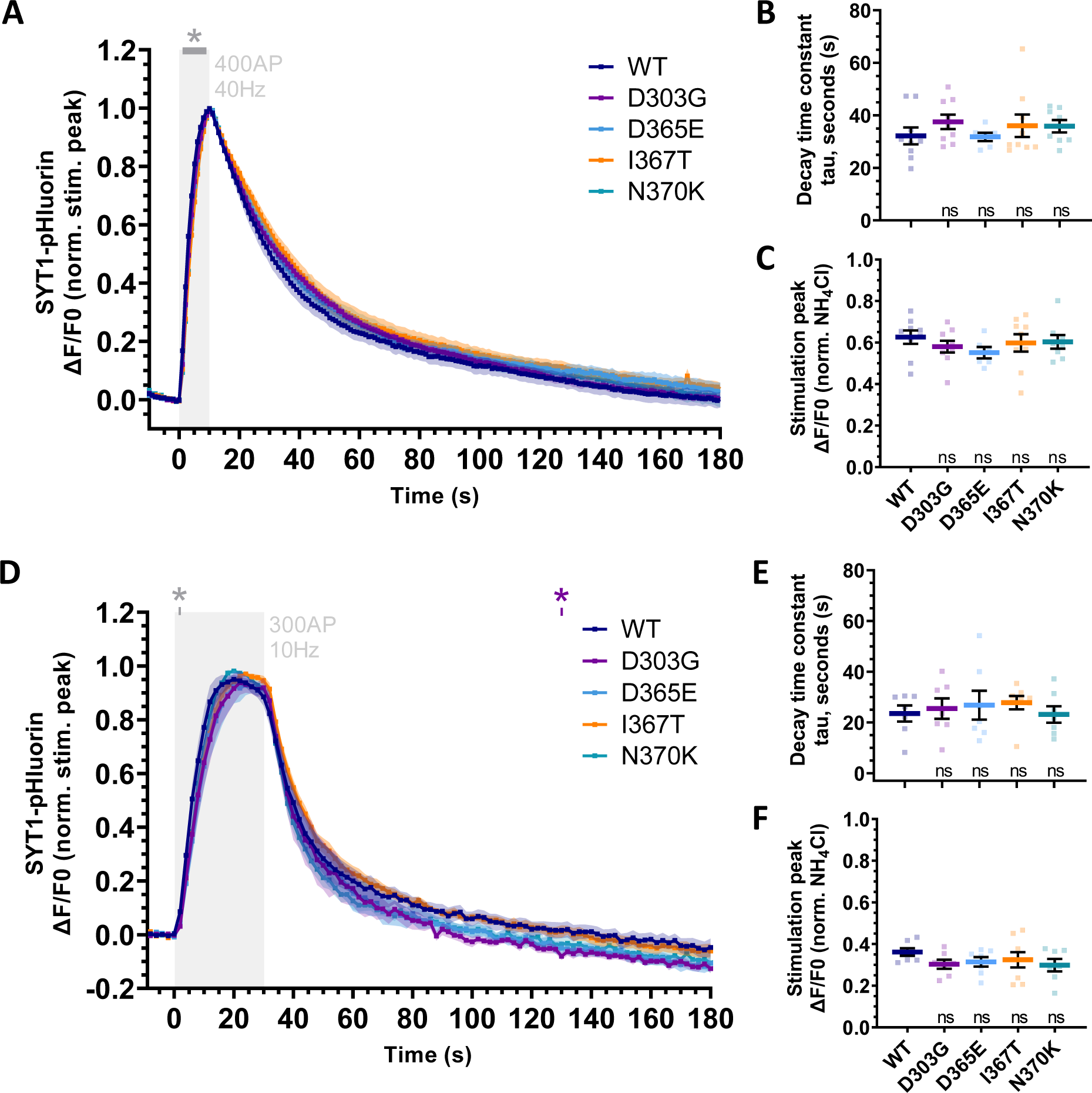
Synaptic vesicle endocytosis is not altered by SYT1 variants. WT neurons were transfected with SYT1-pHluorin variants (WT, D303G, D365E, I367T or N370K) and imaged in the presence of 10 μM CNQX and 50 μM AP5. Baseline images were taken for one minute, before neurons were stimulated either with 400AP at 40 Hz for 10 seconds (A-C; n =6 - 9) or with 300AP at 10 Hz for 30 seconds (D-F; n = 7 - 8). Endocytic retrieval was monitored after the cessation of stimulation, before neurons were superfused with NH_4_Cl buffer to reveal total SYT1-pHluorin fluorescence. Values are mean ± SEM. A,D) Time traces of SYT1-pHluorin fluorescence over the course of the experiments. Fluorescence change from baseline (ΔF/F0) was normalised to the peak fluorescence response to stimulation. A) For 40 Hz for 10 seconds stimulation, two-way repeated measures ANOVA with Dunnett’s multiple comparison’s test was not significant, p>0.05, for all variants compared to WT at all time points, except during stimulation (across 1 to 9 seconds) where *p<0.05 for I367T and N370K (grey asterisk). D) For 10 Hz for 30 seconds stimulation, two-way repeated measures ANOVA with Dunnett’s multiple comparison’s test was not significant, p>0.05, for all variants compared to WT at all time points, except at 2 seconds after stimulation onset where *p<0.05 for D303G, D365E, and I367T (grey asterisk), and at 130 seconds after stimulation onset where *p=0.048 for D303G (purple asterisk). B,E) Time constant (tau) of one-phase exponential decay of SYT1-pHluorin fluorescence following stimulation either at 40 Hz for 10 seconds (B) or at 10 Hz for 30 seconds (E). One-way ANOVA with Dunnett’s multiple comparison’s test was not significant, p>0.05 for all variants compared to WT, for both stimulation paradigms. C,F) Peak SYT1-pHluorin fluorescence response (ΔF/F0) to stimulation either at 40 Hz for 10 seconds (C) or at 10 Hz for 30 seconds (F), normalised to NH_4_Cl maximum SYT1- pHluorin fluorescence. One-way ANOVA with Dunnett’s multiple comparison’s test was not significant, p>0.05 for all variants compared to WT, for both stimulation paradigms. Exact P-values can be found in Table S1.

### Differential impacts of Ca^2+^-binding SYT1 mutants on evoked exocytosis

Having established that endocytosis and spontaneous exocytosis are not affected by SYT1 variants in a dominant-negative manner, disruption to evoked exocytosis therefore remains a major determinant of SYT1 variant pathogenicity. We were consequently interested in examining the molecular mechanisms that contribute to the variation in severity of exocytic impairment and may thereby influence clinical phenotype. Although both variants alter key Ca^2+^-binding aspartate residues of the C2B domain (Figure 4A), D303G and D365E variants result in differing severity of exocytic impairment and neurodevelopmental symptoms, with D365E having a notably milder impact (Baker et al., 2018; Bradberry et al., 2020). We considered whether phenotypic variation arises from the different biochemical properties of the substituting amino acid or from the contributions of different aspartate residues to Ca^2+^ coordination and neurotransmitter release. Additional strategic SYT1 mutants were generated to answer these questions, where the substituted amino acids at each site were swapped, creating D303E and D365G SYT1-pHluorin variants that could be compared to pathogenic variants. These SYT1-pHluorin variants were transfected into cultured WT neurons, and SYT1 immunolabelling confirmed that expression levels and subcellular localisation of mutants were similar to that of WT SYT1-pHluorin (Supplementary Figure 1). For functional assessment, transfected cultures were stimulated at 10Hz for 2 minutes at 37°C degrees in the presence of bafilomycin to evoke exocytosis of the entire recycling pool of synaptic vesicles, and finally superfused with NH_4_Cl buffer (Figure 4A).

**Figure 4.**
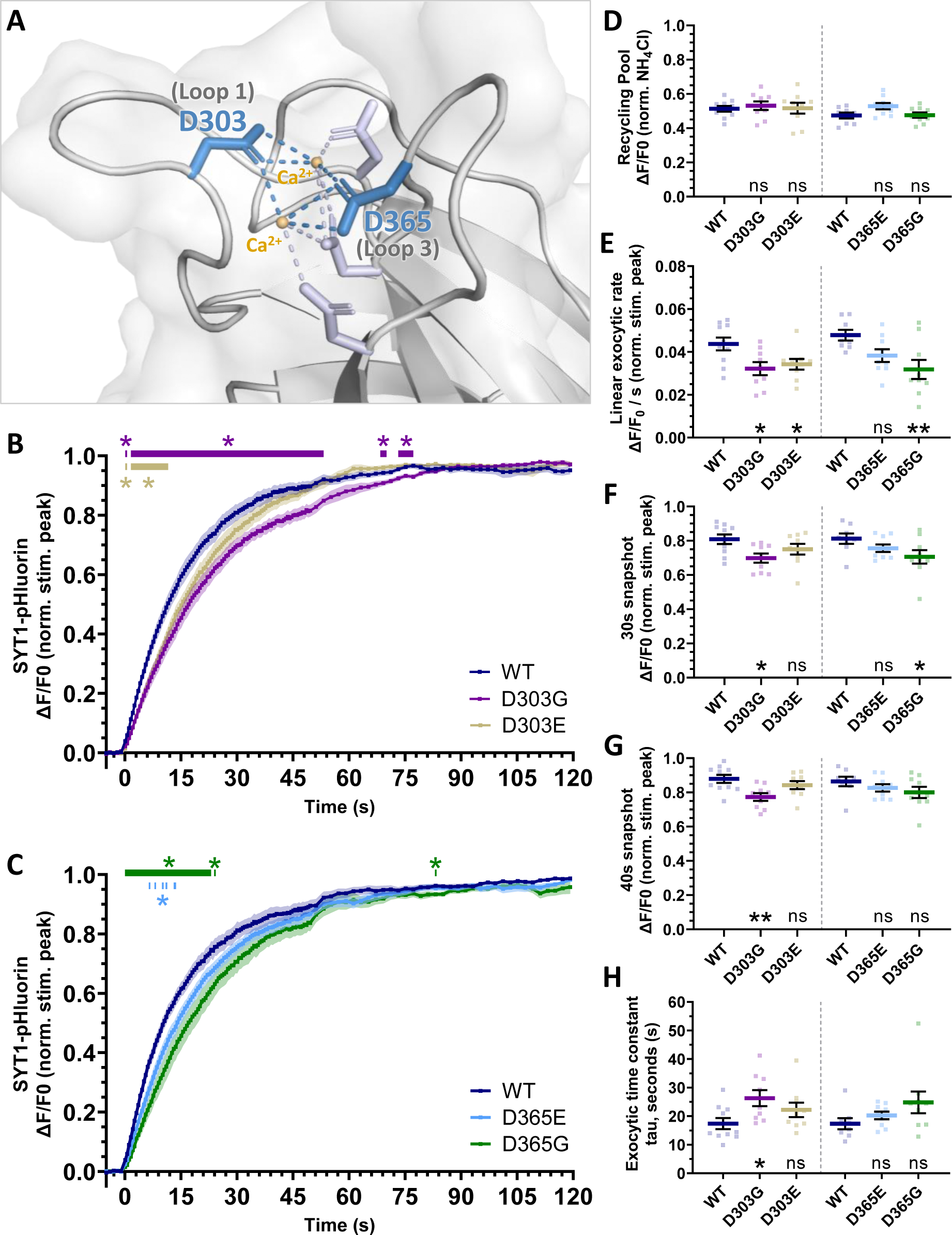
Differential impacts of Ca^2+^-binding SYT1 mutants on evoked exocytosis. WT neurons were transfected with either WT SYT1-pHluorin, pathogenic SYT1-pHluorin variants (D303G, D365E), or strategic SYT1-pHluorin variants (D303E and D365G). Cultures were imaged in the presence of 1 μM bafilomycin A1, 10 μM CNQX, and 50 μM AP5. Cultures were stimulated at 10 Hz for 2 minutes (1200AP) prior to NH_4_Cl superfusion. Fluorescence change from baseline (ΔF/F0) was normalised to peak fluorescence response to stimulation except where otherwise indicated. Values are mean ± SEM, n = 8-10. Variants were compared to their respective WT controls by either two-way repeated measures ANOVA (B,C) or one-way ANOVA (D-H) and with Dunnett’s multiple comparison’s test. A) Three-dimensional structure of the Ca^2+^-binding pocket of the rat synaptotagmin-1 C2B domain (Protein Data Bank: 1K5W), rendered in PyMOL v2.4.0 (Schrödinger, LLC). Ca^2+^ ions are represented as small orange spheres. Ca^2+^-binding aspartate residues are shown in stick representation with the two residues that are impacted by human pathogenic variants and strategically mutated in this study, D303 (D1) and D365 (D4), labelled and coloured blue. Ca^2+^ coordination by aspartate residues is indicated by dashed lines. B,C) Time traces of average SYT1-pHluorin fluorescence over the course of stimulation. Variants significantly differed from WT controls at all time points indicated by horizontal bars, *p<0.05. D) The size of the recycling pool (total vesicle pool mobilised by the stimulus train) was calculated from the fluorescence change from baseline (ΔF/F0) at the end of stimulation (average of last 10 frames) when normalised to peak NH_4_Cl fluorescence and was not significantly different from WT for any variant, p>0.05. E) The linear rate of SYT1-pHluorin fluorescence increase per second over the first 10s of stimulation was significantly slower for D303G, D303E and D365G, but no different for D365E when compared with their respective WT SYT1 controls. **p<0.01, *p<0.05, ns = not significant. F) SYT1-pHluorin fluorescence after 30 seconds of stimulation, reporting cumulative exocytosis evoked by 300AP, was significantly lower for D303G and D365G, but no different for D303E or D365E when compared with their respective WT SYT1 controls. *p<0.05, ns = not significant. G) SYT1-pHluorin fluorescence after 40 seconds of stimulation, reporting cumulative exocytosis evoked by 400 AP, was significantly lower for D303G, but no different for any other variant when compared with their respective WT SYT1 controls. **p<0.01, ns = not significant. H) Time constant (tau, in seconds) of the one-phase exponential increase in SYT1-pHluorin fluorescence during stimulation, was only significantly lower for D303G but no other variant when compared with their respective WT SYT1 controls. *p<0.05, ns = not significant. See Table S1 for exact P-values.

The size of the recycling pool of synaptic vesicles did not differ from WT for any SYT1-pHluorin variant (Figure 4D) but all variants slowed the rate of exocytosis, albeit to differing extents (Figures 4B, 4C and 4E). Pathogenic variants D303G and D365E recapitulated the previously observed mutation-specific severity of impact (Figure 4, (Baker et al., 2018; Bradberry et al., 2020)). At each site, impairment of exocytosis was more severe for glycine substitutions than glutamate substitutions (Figures 4B, 4C and 4F). D303G exerted the greatest impact on exocytosis of all variants (Figures 4B and 4C), being the only variant with significantly lower pHluorin fluorescence (i.e. cumulative exocytic events) than WT SYT-pHluorin after 40 seconds of stimulation (Figure 4G) and with a significantly slower exocytic time constant than the WT control (Figure 4H). D303G can therefore be interpreted to result in more severe effects than D365G. D365E demonstrated the least impact on exocytosis, with a time course of exocytosis that only differed from the WT control at several non-continuous time points; additionally, D365E displayed no significant difference from the WT control in initial linear exocytic rate (Figures 4C and 4E). This is notably milder than the effects of D303E, which resulted in a significantly slower linear exocytic rate over the first 10s of stimulation than WT control (Figures 4B and 4E). 30 seconds after the onset of stimulation, D365G displayed significantly lower fluorescence than WT SYT-pHluorin, whereas D303E did not (Figure 4F). Furthermore, when examining fluorescence time traces, D365G was significantly different from the WT control for a greater number of timepoints than D303E (Figures 4B and 4C). Hence, the relative degree of impairment of exocytosis by each variant can be interpreted as D303G>D365G>D303E>D365E. These strategic swapped mutants therefore reveal that the specific amino acid substitution as well as the site of the mutated aspartate residue contribute to determining the severity of pathogenic exocytic defects, and support that aspartate residues in the Ca^2+^-binding motif of SYT1 may be differentially weighted in their importance for effective Ca^2+^-dependent neurotransmitter release (Nishiki and Augustine, 2004a).

## DISCUSSION

Here, we have delineated the impact of SYT1 variants on distinct aspects of synaptic vesicle dynamics, refining our understanding of the pathogenic mechanisms of SYT1-associated neurodevelopmental disorder. We have established that SYT1 variants primarily exert their pathogenic effects through dominant-negative impairment of evoked exocytosis, with no notable impacts to protein trafficking, proportion of active synapses, spontaneous release, or endocytosis in a disease model context. Moreover, we have revealed that both the identity of the amino acid substitution, as well as the amino acid location, contribute to genotype-phenotype relationships for the Ca^2+^ binding variants.

### SYT1 variants do not have a dominant-negative effect on spontaneous neurotransmitter release

Given that SYT1 clamps spontaneous synaptic vesicle fusion (Broadie et al., 1994; Courtney et al., 2019; Pang et al., 2006; Xu et al., 2009), we sought to ascertain whether increased spontaneous release contributes to the pathogenicity of SYT1 variants. Expression of exogenous SYT1 harbouring patient mutations alongside endogenous WT SYT1 did not result in dominant-negative alterations to the cumulative amount or rate of spontaneous release in our model system, suggesting that disruption to spontaneous synaptic vesicle fusion is not a notable feature of SYT1-associated neurodevelopmental disorder. Further, this supports the theory that clamping of spontaneous release and facilitation of evoked synchronous release are underpinned by distinct molecular mechanisms involving SYT1 (Kaeser and Regehr, 2014; Kavalali, 2015).

Our results contrast with a previous study observing that the D303G SYT1 variant does not restore the clamping of spontaneous release at inhibitory synapses when expressed on a SYT1 null background (Bradberry et al., 2020). This suggests that the presence of WT SYT1 is sufficient to fulfill this clamping function. Few studies have specifically examined whether mutations in the SYT1 Ca^2+^-binding loops can impact spontaneous release in a dominant-negative manner. C2B Ca^2+^-binding loop mutants have shown mixed results in Drosophila larval neuromuscular junction preparations also expressing WT SYT1 (Guan et al., 2017; Herrmann et al., 2014; Lee et al., 2013; Shields et al., 2017), but these studies are confounded by the fact that the molecular players controlling spontaneous fusion in Drosophila differ from those in mammalian central synapses (Courtney et al., 2019; Sauvola and Littleton, 2021; Trimbuch and Rosenmund, 2016; Xu et al., 2009). A SYT1 triple mutant (“3DA”: D309A/D363A/D365A) showed a dominant-negative increase in spontaneous release when expressed in mammalian cultured neurons (Zhou et al., 2017); however, this highly deleterious triple mutant is expected to abolish Ca^2+^ binding to the C2B domain and is not reflective of known pathogenic variants (Shin et al., 2009).

Interestingly, a neurodevelopmental disorder-associated SYT1 variant, P401L (equating to P400L in mouse/rat sequence), was recently demonstrated to increase spontaneous miniature excitatory postsynaptic currents in a dominant-negative manner in autaptic murine neurons (Cornelisse et al., 2023). This variant is notably situated on the opposite side of the C2B domain from the Ca^2+^-binding loop, lying adjacent to the functionally important arginine apex. It also showed a distinct cellular phenotype where the inhibitory role of SYT1 in clamping asynchronous and spontaneous release was impaired, but not the ability of SYT1 to promote action-potential evoked synaptic vesicle fusion, which contrasts the established mechanisms of the Ca^2+^-binding loop variants (Baker et al., 2018; Bradberry et al., 2020). Additionally, a strategic variant located on the distal end of the C2B domain from the Ca^2+^-binding loops, F349A, which impairs SYT1 oligomerisation, also caused a dominant-negative increase in mEPSC frequency when introduced into wild-type murine cultures (Bello et al., 2018; Tagliatti et al., 2020). The role of the C2B domain of SYT1 in regulating spontaneous neurotransmitter release is evidently nuanced and the specific residues involved in this function remain enigmatic. Beyond this, further investigation is still required to understand the specific mechanisms governing spontaneous and evoked synaptic vesicle fusion at mammalian synapses and the molecular interplay between mutant and WT SYT1.

### Disease-associated SYT1 variants do not affect endocytosis

SYT1 variants also did not alter the rate of endocytosis in a dominant-negative manner in response to varied frequencies and durations of stimuli, indicating that the pathophysiology of this disorder does not involve perturbation of synaptic vesicle endocytosis. Previously we showed that D303G and D365E SYT1 remain diffusely localised following strong neuronal depolarisation, suggesting delayed retrieval to nascent synaptic vesicles; however, this was in response to non-physiological KCl stimulation (Baker et al., 2018). Additionally, I367T SYT1 was initially shown to increase the rate of endocytosis when assayed at room temperature (Baker et al., 2015). However, the kinetics of endocytosis are highly sensitive to temperature and the prevalence of distinct modes of endocytosis differ between room and physiological temperatures (Chanaday et al., 2019; Chanaday and Kavalali, 2018; Gan and Watanabe, 2018; Watanabe and Boucrot, 2017). Electrical stimulation at physiological temperature, as was performed in the present study, is therefore a better model for synaptic vesicle recycling in the human brain, and highlights the importance of using physiologically relevant stimuli and temperatures when assaying presynaptic activity.

The roles of the disease-relevant SYT1 residues in endocytosis have otherwise only been investigated in combination with other mutated amino acids and when expressed on SYT1 KO background. A double D363,365N mutant (or equivalent in drosophila) has been examined in two separate studies: one demonstrating a slowing of endocytosis at drosophila larval neuromuscular junctions (Poskanzer et al., 2006), and the other showing no difference from WT SYT1 in cultured murine neurons (Yao et al., 2011). Additionally, mutating all four membrane penetrating hydrophobic residues at the tips of the Ca^2+^-binding loops of the C2A and C2B domains (M173, F234, V304 and I367) to either alanine or tryptophan did not affect the rate of endocytosis (Yao et al., 2011). Collectively, this supports that neurodevelopmental disorder-associated SYT1 C2B domain variants are unlikely to impact the function of SYT1 as a modulator of endocytosis.

### Ca^2+^-binding mechanisms of the SYT1 C2B domain

We next addressed the differing clinical and cellular phenotypic severity of two SYT1 variants, D303G and D365E. Is phenotypic variability due to substitution with distinct amino acids or mutation of different Ca^2+^-binding sites? Comparing human mutations with strategic mutation of the same residues, we found that both the site of the mutated residue and the specific amino acid substitution contribute to the severity of impact on evoked exocytosis. The divergence in symptom severity observed between individuals harbouring D303G and D365E variants can therefore likely be attributed to a combination of these influences.

This result is supported by previous findings that aspartate residues in the C2B Ca^2+^-binding motif are differentially weighted in their contribution to facilitating evoked exocytosis. While usually studied in tandem, Nishiki and Augustine (2004a) mutated each aspartate to asparagine individually, revealing that D2 and D3 mutants (D309 and D363 in rat) failed to rescue evoked neurotransmitter release and the D5 mutant (D371) fully rescued release, when expressed in SYT1 KO neuronal cultures. D1 and D4 (D303 and D365) mutants had intermediate impacts; both impaired neurotransmitter release, though D365N to a lesser extent than D303N (Nishiki and Augustine, 2004a). We observed this same graded contribution with both glutamate and glycine substitutions at D303 and D365 (Figure 4), therefore Ca^2+^ binding to D303 is likely of greater importance for the effectiveness of evoked synaptic vesicle fusion than Ca^2+^ binding to D365.

However, it is clearly important to also consider the specific amino acid change. Many studies have examined only asparagine or alanine substitutions of Ca^2+^-binding aspartates. It has been suggested that an aspartate to asparagine neutralising mutation partially mimics the binding of Ca^2+^, whereas substituting with glutamate would conserve the charge at that site but may hinder Ca^2+^- binding, potentially resulting in substantially different impacts (Shields et al., 2020; Striegel et al., 2012). Our work showed that glycine substitution resulted in more severe exocytic defects than glutamate substitution at both D303 and D365 sites (Figure 4). The pathogenicity of each new disease-linked SYT1 variant should be considered carefully as different substitutions at the same site could have markedly different impacts on SYT1 function and consequently phenotype; caution is therefore needed when predicting clinical prognosis involving novel substitutions at recurrent loci. Additionally, our results reinforce that even conservative substitutions, such as aspartate to glutamate, can impair neurotransmitter release when they affect highly sensitive residues such as the Ca^2+^-binding aspartates of the C2A and C2B domains (Baker et al., 2018; Bradberry et al., 2020; Shields et al., 2020; Striegel et al., 2012). Careful consideration and functional characterisation of emerging disease-associated variants may improve prognostication and provide insight into the exact molecular mechanisms underpinning SYT1 function. Additionally, dissection of the calcium binding mechanisms of SYT1 will further benefit from the use of other strategic single residue mutations to probe the precise electrostatic interactions involved in the regulation of neurotransmitter release by SYT1.

Additional variants in SYT1 have recently been reported, many of which are situated outside of the Ca^2+^-binding pockets, with several located in the C2A domain (Melland et al., 2022). For some of these variants, the single cases reported to date are associated with a clinical phenotype that is notably milder than for any variants examined in the present study (i.e. milder developmental delay and intellectual disability, no movement disorder). Functional characterisation of new SYT1 variants is required to confirm whether these newly-identified variants also exert their pathogenic effects through impairment of evoked synaptic vesicle exocytosis (though possibly to a lesser degree than variants examined here) or rather though altering other aspects of synaptic vesicle dynamics. For example, phenotypic heterogeneity within another disorder of synaptic vesicle fusion has been linked to distinct mechanistic aetiologies, where variants in *SNAP25* differentially impacted properties of evoked and spontaneous neurotransmitter release (Alten et al., 2021). Together with previous work on Ca^2+^-binding loop variants in SYT1 (Baker et al., 2015; Baker et al., 2018; Bradberry et al., 2020), one recent study of a P401L (human) variant in SYT1 (Cornelisse et al., 2023) supports that such mechanistic heterogeneity may also be present within SYT1-associated neurodevelopmental disorder.

Nevertheless, current evidence from four extensively characterised, pathogenic SYT1 variants suggests that impairments in evoked exocytosis is a central presynaptic pathological feature of SYT1- associated neurodevelopmental disorder. As such, we confirm that this syndrome can accurately be considered a “disorder of synaptic vesicle fusion”, an emerging group of neurological diseases arising from various disruptions to synaptic vesicle exocytosis (Melland et al., 2021). This delineation of the precise pathophysiology of SYT1-associated neurodevelopmental disorder further clarifies mechanistic targets for treatment and could improve confidence in predicting and preventing off-target therapeutic effects.

## MATERIALS AND METHODS

### Reagents

QuikChange II Site-Directed Mutagenesis Kit and Dako Fluorescence Mounting Medium were from Agilent Technologies (Santa Clara, CA, USA). Neurobasal Medium, Dulbecco’s Modified Eagle Medium: Nutrient mixture F-12 (DMEM/F12), Minimum Essential Medium with Earle’s salts and L-glutamine (MEM), penicillin/streptomycin, L-glutamine, B-27 Supplement 50x (B-27), Lipofectamine 2000, goat anti-chicken IgY (H+L) Alexa Fluor 488, goat anti-chicken IgY (H+L) DyLight 550, and donkey anti-rabbit IgG (H+L) Alexa Fluor 647 were obtained from ThermoFisher Scientific (Scoresby, Australia). Rabbit anti-SYT1 was from Synaptic Systems (Göttingen, Germany). Chicken anti-GFP was from Abcam (Melbourne, Australia). Papain was sourced from Worthington Biochemical Corporation (Lakewood, NJ, USA). HyClone Fetal Bovine Serum (FBS) was from In Vitro (Melbourne, Australia). Coverslips were sourced from VWR International (Radnor, PA, USA). 6-cyano-7-nitroquinoxaline-2,3-dione (CNQX), DL- 2-Amino-5-phosphonopentanoic acid (AP5), and Bafilomycin A1 were distributed by Sapphire Bioscience (Sydney, Australia) from Enzo Life Sciences (Lausen, Switzerland), Cayman Chemical (Ann Arbor, MI, USA) and Toronto Research Chemicals (Toronto, Canada), respectively. All other reagents were obtained from Sigma-Aldrich (Castle Hill, Australia).

### DNA constructs and mutagenesis

SYT1-pHluorin and mCherryN1 constructs were kindly provided by Professor V. Haucke (Leibniz Institute of Molecular Pharmacology, Berlin, Germany) and Professor M. A. Cousin (University of Edinburgh, UK), respectively. SYT1-pHluorin D303G, D365E, I367T and N370K mutant constructs were generated as described in Baker 2015 and Baker 2018. Mutagenesis of SYT1-pHluorin to induce D303E and D365G variants was performed using QuikChange II Site-directed Mutagenesis kit, using either the included PfuUltra High-Fidelity DNA Polymerase (D365G) or KOD Hot Start Polymerase (D303E), and primers (D303E: forward – CCTGAAGAAGATGGAGGTGGGTGGCTTATACTGA, reverse – GATCAGATAAGCCACCCAC**<u>C</u>**TCCATCTTCTTCAGG; D365G: forward – GTTTTGGACTATG**<u>G</u>**CAAGATTGGCAAGAACGACGC, reverse - GCGTCGTTCTTGCCAATCTTG**<u>C</u>**CATAGTCCAAAAC . Mutations were verified by DNA sequencing (performed by the Australian Genome Research Facility, Melbourne, Australia).

### Primary hippocampal neuronal culture

All procedures were approved by the Florey Animal Ethics Committee and performed in accordance with the guidelines of the National Health and Medical Research Council Code of Practice for the Care and Use of Animals for Experimental Purposes in Australia. Mouse colonies were maintained in a temperature controlled (≈ 21°C) room and group housed in individually ventilated cages on a 12 hour light-dark cycle (lights on 07:00–19:00) with food and water available ad libitum. Animals were time mated overnight and visualisation of a vaginal plug on the following morning was considered as embryonic day (E) 0.5.

Dissociated primary hippocampal-enriched neuronal cultures were prepared from hippocampi dissected from E16.5-18.5 C57BL/6J mouse embryos of both sexes. Hippocampi were incubated in 10 units/mL papain in PBS at 37°C for 20 minutes. Papain was removed and replaced with DMEM/F12 supplemented with 10% v/v FBS and 1% v/v penicillin/streptomycin before cells were mechanically disaggregated into a single cell suspension by careful trituration. Cells were pelleted by centrifugation at 363g for 5 minutes, DMEM/F12 supernatant was replaced with Neurobasal media, and cells were resuspended prior to plating. Cells were plated on poly-D-lysine and laminin-coated coverslips at a density of approximately 3.5x10^4^ cells per 13mm coverslip in 24 well plates for fixation and immunolabelling or at a density of approximately 5x10^4^ cells per 25mm coverslip in 6 well plates for live cell imaging. Cultures were maintained in a humidified incubator at 37°C and 5% CO_2_ in Neurobasal media supplemented with B-27, 0.5mM L-glutamine and 1% v/v penicillin/streptomycin. After 48-96 hours, cultures were further supplemented with 1μM cytosine β-d-arabinofuranoside to inhibit glial proliferation. Cells were transfected after 6-8 days in culture with SYT1-pHluorin variants alongside either an empty pcDNA3.1/Zeo vector for fixation and immunolabelling experiments or mCherryN1 (as a transfection marker) for live cell imaging. Lipofectamine 2000 transfection agent was used according to the manufacturer’s instructions with the following alterations: cells were incubated for 2 hours at 37°C with 5% CO_2_ in MEM with 1 μL Lipofectamine 2000 and 0.5 μg of each DNA construct per well for 24 well plates (for fixation and immunolabelling) or 2 μL of Lipofectamine 2000 and 1 μg of each DNA construct per well for 6 well plates (for live cell imaging). Cells were subsequently washed with MEM before replacement of conditioned Neurobasal media. Cells were either fixed or used for live cell imaging assays after 13-16 days in culture.

### Live Cell Imaging and Analysis

Live neuronal cultures were mounted in a Warner imaging chamber with embedded parallel platinum wires (RC-21BRFS). Electrical field stimulation was applied using a Digitimer D330MultiStim System from SDR Scientific (Chatswood, Australia) at 35 mA with 1 ms pulse width. All neurons were constantly superfused with saline buffers at 37°C using a Warner TC-324C in-line heater. All buffers were superfused for at least one minute before fluorescence measurements were taken.

Analysis of time series images was undertaken using the FIJI 1.52n distribution of ImageJ. Experimenters were blinded to the SYT1-pHluorin variant for image and data analysis. To correct for slight XY-plane drift over the time series, stack registration was performed by rigid body alignment to the first frame of the experiment using the MBGReg ImageJ plugin (Donal Stewart, Edinburgh, UK), a custom variant of the FIJI StackReg plugin. To remove the component of fluorescence decay contributed by the photobleaching of background autofluorescence in time series images from endocytosis) and evoked exocytosis assays, the Bleach Correction Image J plugin was used (version: CorrectBleach_-2.0.3-SNAPSHOT.jar, https://github.com/fiji/CorrectBleach) (Miura, 2020). Photobleaching over the full length of the experiment was corrected using a single exponential function fitted to the fluorescence decay of background regions of interest (ROIs) over the duration of the experiment prior to NH_4_Cl buffer superfusion.

Identically-sized circular ROIs were manually placed around presynaptic boutons (fluorescence puncta) using the Time Series Analyzer V3 plugin (https://imagej.nih.gov/ij/plugins/time-series.html), and the total pHluorin fluorescence intensities of each ROI were calculated for each frame of the time series. Fluorescence time traces of each ROI were manually screened for bona fide responses using an in-house screening tool (Holly Melland). Only ROIs that increased fluorescence from baseline upon both electrical stimulation and NH4Cl buffer superfusion were included in subsequent analysis. In Microsoft Excel, fluorescence responses of each ROI were calculated as ΔF/F0 (change in fluorescence from baseline). For each frame of the time series, fluorescence values were averaged for all ROIs in a field of view, except where otherwise specified.

### Synaptic vesicle targeting assay

Neurons were imaged on a Leica TCS SP8 inverted scanning confocal microscope inside an incubation chamber maintained at 37°C and 5% CO_2_. SYT1-pHluorin variants were visualised through a Leica HC PL APO CS2 40x (NA 1.3) oil-immersion objective with an Argon laser 488 nm excitation wavelength and 501-556 nm emission window. Single channel, 16-bit images of SYT1-pHluorin fluorescence were acquired using a hybrid photodetector with a pixel size of 0.284 μm. Confocal Z-stacks including the entire volume of the presynaptic terminals of interest were acquired and Nyquist sampling was used to calculate a z-step size of 0.3 μm. Image analysis was performed on the summed projection of Z-slices. Z-stack images were acquired every 30 seconds and neuronal cultures were successively superfused with different imaging buffers to probe SYT1-pHluorin in distinct membrane compartments. Standard saline imaging buffer (136mM NaCl, 2.5mM KCl, 2mM CaCl2, 1.3mM MgCl2, 10mM glucose, 10mM HEPES, pH 7.4) obtained a baseline fluorescence value reflecting the surface population of SYT1-pHluorin, an acidic imaging buffer (20 mM MES in place of HEPES, pH 5.5) quenched surface pHluorin fluorescence and yielding a background value, and an alkaline NH_4_Cl buffer (50 mM NH_4_Cl substituted for 50 mM NaCl, pH 7.4) exposed total SYT1-pHluorin fluorescence. The percentage of internalised SYT1-pHluorin, indicating SYT1-pHluorin targeted to synaptic vesicles, was calculated as 100 - 100x((F baseline – F acid)/(F NH_4_Cl – F acid)).

### Exocytosis and endocytosis assays

Neurons were imaged on a Zeiss Axio Observer 7 inverted epifluorescence microscope with a Colibri 7 LED light source. Cells were visualised through a Zeiss EC Plan-Neofluar 40x oil-immersion objective (NA 1.3), and a Zeiss Axiocam 506 mono camera captured 14-bit images with 2x2 binning yielding a pixel size of 0.227 µm. SYT1-pHluorin was visualised with a 475 nm excitation wavelength through a single band pass GFP filter set (excitation 470/40, emission 525/50, beam splitter 495 nm). For the spontaneous exocytosis assay, mCherry was visualised with a 567 nm excitation wavelength through a single bandpass DsRed filter set (excitation 550/25, emission 605/70, beam splitter 570 nm). Within each experiment, all images were acquired with identical illumination intensity and exposure length. Automated triggering of electrical stimulation and variable image acquisition rate was implemented using the Zen Blue Experiment Designer module. Neurons were superfused with NH_4_Cl buffer at the end of each assay to reveal total SYT1-pHluorin fluorescence.

For the spontaneous exocytosis assay, baseline fluorescence was captured for one minute while neurons were superfused with standard saline imaging buffer. Cultures were then superfused with standard saline imaging buffer supplemented with 1 μM bafilomycin A1, 10 μM CNQX, and 50 μM AP5. Spontaneous release was captured with one image every 30 seconds for 10 minutes. Evoked release was elicited through electrical stimulation at 10 Hz for 30 seconds (300 AP) with images captured at 0.5Hz.

For the endocytosis assay, neuronal cultures were superfused with standard saline imaging buffer supplemented with 10 μM CNQX and 50 μM AP5. Images of SYT1-pHluorin fluorescence were acquired at 1 or 2Hz. Neurons were electrically stimulated at 10Hz for 30s (300AP) or 40 Hz for 10 seconds (400 AP), and subsequent endocytic retrieval of SYT1-pHluorin was observed for 150 or 170 seconds.

For the evoked exocytosis assay, cultures were superfused with standard saline imaging buffer supplemented with 1 μM bafilomycin A1, 10 μM CNQX, and 50 μM AP5, and electrically stimulated at 10 Hz for 2 minutes (1200 AP), with images captured at 2 Hz for 50 seconds then 0.5 Hz for 70 seconds.

### Statistical Analysis

Experimenters were blinded to the SYT1-pHluorin variant for image analysis. Data were compiled, calculated and averaged in Microsoft Excel and appropriate statistical analyses were performed in GraphPad Prism v8 and v9. All data were tested for normality prior to parametric testing. All data are presented as mean ± standard error of the mean (SEM). Descriptive statistics and exact P-values are listed in Supplementary Table 1. Each n represents a single field of view from an independent coverslip, and all experiments were performed across at least 3 independent cultures.

## ACKNOWLEDGEMENTS AND FUNDING

We appreciate the important and generous contributions of each individual with a SYT1 variant, their families, and carers. We appreciate the roles of each clinician and laboratory scientist involved in the diagnostic pathway of each case. We thank the Core Animal Services staff and Bioresources Facilities of the Florey Institute for animal maintenance and breeding. We appreciate past and present Gordon laboratory members for their useful discussions. This work was supported by National Health and Medical Research Council (NHMRC) Ideas Grant (2003710) and a Florey Fellowship to S.G.. H.M. and E.H.A. were supported by an Australian Government Research Training Program Scholarship. K.B. is supported by the UK Medical Research Council (G116768). The Florey Institute of Neuroscience and Mental Health acknowledges the strong support from the Victorian Government and in particular the funding from the Operational Infrastructure Support Grant.

## COMPETING INTERESTS

The authors declare no competing financial interests.

## AUTHOR CONTRIBUTION

Conceptualisation: S.L.G and H.M.; Data Curation: H.M.; Formal Analysis: H.M., K.V., S.L.G.; Funding Acquisition: S.L.G.; Investigation: H.M., K.V., S.L.L, A.F. E.H.A; Visualisation: H.M.; Writing-original draft: H.M. (lead), S.G.; Writing-review and editing: H.M., S.L.G., K.B., All other authors reviewed and approved this manuscript.

## DATA AVAILABILITY

Summary data of statistical testing is available in Supplementary Table 1. Raw data are available upon request to the corresponding author.

## Notes

### Competing Interest Statement

The authors have declared no competing interest.

