## Supplementary Material for "Delineation of the pathogenic presynaptic mechanisms of synaptotagmin-1 variants"

### SUPPLEMENTARY FIGURES

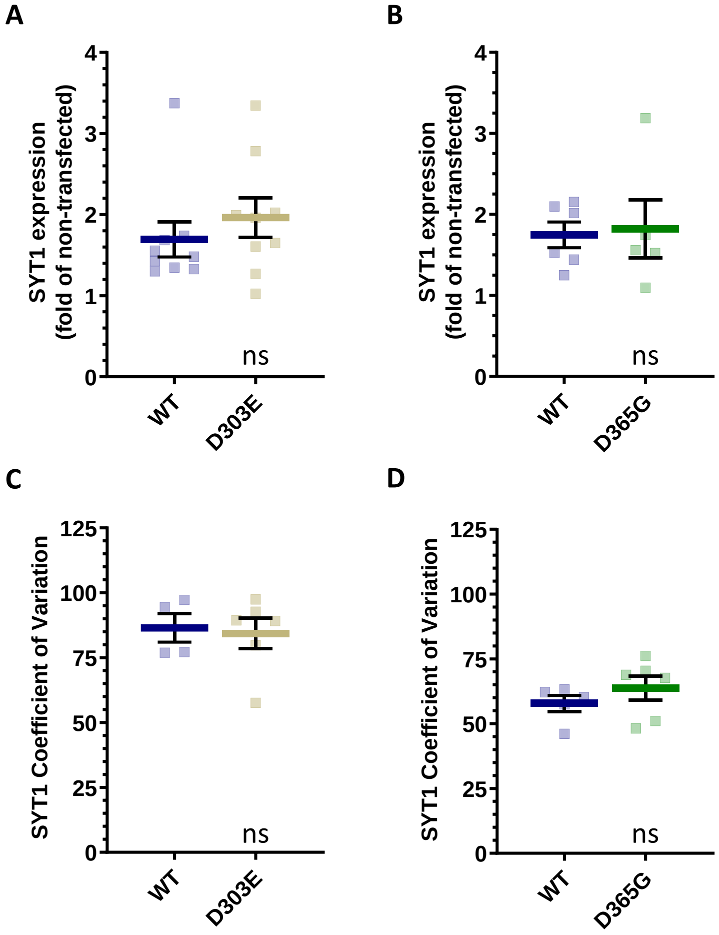


**Supplementary Figure 1. Expression and localisation of flipped SYT1-pHluorin aspartate variants are similar to the WT protein.** Cultured hippocampal neurons were transfected with SYT1 variants and immunolabelled for GFP/pHluorin and SYT1. A-B) SYT1 immunofluorescence intensity at synaptic puncta in transfected neurons relative to non-transfected neurons in the same field of view. A) D303E SYT1-pHluorin expression level. N=9 individual fields of view from independent coverslips across two separate cultures. B) D365G SYT1-pHluorin expression level. N=5-6 individual fields of view from independent coverslips across two separate cultures. C-D) Targeting of SYT1-pHluorin variants to presynaptic terminals was determined by the coefficient of variation of the distribution of SYT1-pHluorin fluorescence intensity along neurites, where a higher coefficient of variation equates to a more punctate localisation. C) Coefficient of variation of D303E SYT1-pHluorin. N=4-6 individual fields of view from independent coverslips across two separate cultures. D) Coefficient of variation of D365G SYT1-pHluorin. N=5-6 individual fields of view from independent coverslips across two separate cultures. For all panels, P>0.05 compared to WT by unpaired t-test (see Table S1 for exact P-values). ns = not significant.

### SUPPLEMENTARY METHODS

#### Expression and localisation assays

Primary hippocampal neuronal cultures were first washed with standard saline imaging buffer and then fixed in 4% paraformaldehyde in phosphate-buffered saline (PBS) for 20 minutes. Cells were then incubated at room temperature in 50 mM $NH_4Cl$ in PBS for 10 minutes, permeabilised with 0.1% v/v Triton X-100, 1% v/v bovine serum albumin (BSA) in PBS for 5 minutes, and blocked with 1% BSA in PBS for at least one hour before immunolabelling, with PBS washing between each step.

Cells were incubated in primary and then secondary antibodies in 1% BSA in PBS for 1-2 hours (with extensive washing with PBS after each incubation), washed in Milli-Q $H_20$ and mounted on glass slides with Dako Fluorescence Mounting Medium. Primary antibodies used were chicken anti-GFP (recognising pHluorin; 1:2000, Abcam, ab13970) and rabbit anti-SYT1 (1:100, SYSY, 105102). Fluorescent secondary antibodies used were Alexa Fluor 488 goat anti-chicken (1:500, Thermo, A11039) or Dylight 550 goat anti-chicken (1:500, Thermo, SA5-10071) and Alexa Fluor 647 donkey anti-rabbit (1:500, Thermo, A31573).

D303E experiments were imaged on a Zeiss Axio Observer.Z1 inverted epifluorescence microscope with HXP 120 V mercury arc lamp. Cells were visualised through a Zeiss Plan-Apochromat 63x oil-immersion objective (NA 1.4) and a Zeiss Axiocam 506 mono camera captured 14-bit images with 2x2 binning yielding a pixel size of 0.144 µm. DyLight 550-conjugated anti-chicken (recognising anti-GFP) and Alexa Fluor 647-conjugated anti-rabbit (recognising anti-SYT1) secondary antibodies were used. DyLight 550 was visualised through a single bandpass DsRed filter set (excitation 550/25, emission 605/70, beam splitter 570 nm). Alexa Fluor 647 was visualised through a single band pass Cy5 filter set (excitation 640/30, emission 690/50, beam splitter 660 nm).

D365G experiments were imaged on a Zeiss Axio Observer 7 inverted epifluorescence microscope with a Colibri 7 LED light source. Cells were visualised through a Zeiss EC Plan-Neofluar 40x oil-immersion objective (NA 1.3), and a Zeiss Axiocam 506 mono camera captured 14-bit images with 2x2 binning yielding a pixel size of 0.227 µm. To assess the expression of SYT1-pHluorin D365G, Alexa Fluor 488-conjugated anti-chicken (recognising anti-GFP) and Alexa Fluor 647-conjugated anti-rabbit (recognising anti-SYT1) secondary antibodies were used. To assess the localisation of SYT1-pHluorin D365G, DyLight 550-conjugated anti-chicken (recognising anti-GFP) secondary antibodies were used. Alexa Fluor 488 was visualised with a 475 nm excitation wavelength through a single band pass GFP filter set (excitation 470/40, emission 525/50, beam splitter 495 nm). DyLight550 and Alexa Fluor 647 were visualised with 567 nm and 630 nm excitation wavelengths, respectively, through a quadruple bandpass filter set including Cy3 and Cy5 (excitation 385/30+469/38+555/30+631/33, emission 425/30+514/30+592/25+709/100, beam splitters 405+493+575+653).

Expression and localisation of wild-type (WT), D303E and D365G SYT1-pHluorin variants were analysed using the FIJI 1.52n distribution of ImageJ and the Time Series Analyzer V3 plugin. Experimenters were blinded to the SYT1-pHluorin variant for image acquisition and analysis.

To determine SYT1-pHluorin expression levels, circular ROIs of uniform size were manually placed around synaptic boutons (i.e. punctate SYT1 immunofluorescence) of transfected (GFP/pHluorin immunofluorescence) and non-transfected neurons, as well as in background regions. The total fluorescence intensity was calculated for each ROI. For each field of view, SYT1 expression level in transfected neurons was calculated using Microsoft Excel as fold-change over the non-transfected endogenous SYT1 expression level by dividing the average fluorescence intensity of transfected puncta by that of non-transfected puncta after subtracting background intensity. A single n represents a single field of view.

The localisation of SYT1-pHluorin variants was measured by coefficient of variation analysis. ROIs in the form of freehand lines were drawn along sections of transfected neurite spanning greater than 40 µm and fluorescence intensity values along the length were obtained. In Microsoft Excel, the coefficient of variation was calculated by dividing the standard deviation of the fluorescence values composing each ROI by their mean and multiplying by 100. A single n represents a single field of view, where the coefficient of variation of five lengths of neurite were averaged. A high coefficient of variation is indicative of heterogenous/punctate SYT1-pHluorin immunofluorescence along the neurite (such as is seen for enrichment at *en passant* synaptic boutons), while a low coefficient of variation indicates homogenous/diffuse SYT1-pHluorin immunofluorescence (indicating inefficient trafficking of SYT1, which is normally enriched at synaptic boutons).
